# Mimicry is associated with procedural learning, not social bonding, neural systems in autism

**DOI:** 10.1101/2020.09.16.299370

**Authors:** Bahar Tunçgenç, Carolyn Koch, Amira Herstic, Inge-Marie Eigsti, Stewart Mostofsky

## Abstract

Mimicry facilitates social bonding throughout the lifespan. Mimicry impairments in autism spectrum conditions (ASC) are widely reported, including differentiation of the brain networks associated with its social bonding and learning functions. This study examined associations between volumes of brain regions associated with *social bonding* versus *procedural skill learning*, and mimicry of gestures during a naturalistic interaction in ASC and neurotypical (NT) children. Consistent with predictions, results revealed reduced mimicry in ASC relative to the NT children. Mimicry frequency was negatively associated with autism symptom severity. Mimicry was predicted predominantly by the volume of procedural skill learning regions in ASC, and by bonding regions in NT. Further, bonding regions contributed significantly less to mimicry in ASC than in NT, while the contribution of learning regions was not different across groups. These findings suggest that associating mimicry with skill learning, rather than social bonding, may partially explain observed communication difficulties in ASC.

## Introduction

During a conversation or while watching a movie, people unconsciously copy the facial expressions, postures, and body movements of those they interact with. Copying mannerisms and expressions, from here on referred to as mimicry, is different from copying goal-directed actions, such as imitating the way an expert holds the racket in a tennis game. While imitating goal-directed actions can help with learning new skills, mimicry of mannerisms or body movements can help forge social bonds (Over & Carpenter, 2013). Despite known deficits in both goal-directed and social mimicry in autism spectrum conditions (ASC), the distinct social bonding function of mimicry in relation to brain structures has not been investigated in this population. In this study, we used a naturalistic story-telling task to examine the neural substrates of these divergent functions (i.e., learning versus bonding) in children with ASC and those with neurotypical (NT) developmental histories.

### Mimicry behaviour and autism

Research in NT children and adults has shown that both mimicking and being mimicked are strongly linked with increased interaction quality, social bonding between the interactants, and pro-social behaviours (Duffy & Chartrand, 2015; Stel, van Dijk, & van Baaren, 2016). While mimicry is a prevalent feature of social interactions between NT individuals, it may not be so for individuals with ASC. Impairments in spontaneous mimicry of facial expressions (Oberman, Winkielman, & Ramachandran, 2009), yawning (Helt, Eigsti, Snyder, & Fein, 2010; Senju et al., 2007) and body movements (K. L. Marsh et al., 2013) have been widely reported in ASC. While these impairments often involve reduced mimicry in ASC, a recent study (Helt et al., 2020) has shown that increased mimicry can also be related to autism severity, especially when the actions involve negatively-valenced stimuli that may evoke personal distress (e.g., contagious itching as if sand is in one’s hair or clothes). Studies also show robust autism-associated impairments in copying the style of an action (Hobson & Hobson, 2008) or copying actions that are irrelevant to attaining a goal, such as tapping the lid of a box before opening it (Marsh, Pearson, Ropar, & Hamilton, 2013). Notably, imitation of seemingly meaningless gestures is reduced in children with ASC even when they are explicitly instructed to copy the observed sequences (McAuliffe et al., 2019)(McAuliffe et al., 2020).

This prior work points to an imitation mechanism in ASC, whereby kinematic cues carrying stylistic information are omitted, leading to poorer copying of meaningless, or non-goal-directed, gestures (Gowen, 2012). Such mimicry of seemingly meaningless mannerisms or expressions plays a big role in fostering bonding during typical social interactions (Duffy & Chartrand, 2015). Moreover, being mimicked can improve social gaze, social touch, pointing and play skills in children with autism (Contaldo, Colombi, Narzisi, & Muratori, 2016). Thus, autism-associated mimicry deficits may partially explain observed social-communication difficulties in ASC. Demonstrating this link in ASC children aged 2 to 4 years old, a recent study (Ingersoll & Meyer, 2011) found that spontaneous, but not instructed, imitation was associated with social reciprocity and symbolic play skills, as measured by the standardised autism assessment tool, the Autism Diagnostic Observation Schedule (ADOS).

Building upon this body of literature, we hypothesised that, during a semi-structured social interaction, children with ASC would mimic their partner’s gestures (i.e., face rubbing, arm scratching) less than NT children (Hypothesis 1). Further, we predicted that the degree of mimicry would be negatively associated with autism symptom severity (Hypothesis 2).

### Neural mechanisms of mimicry and autism

To understand the mechanisms underlying communicative difficulties in autism, it is critical to establish whether mimicry serves a social bonding function in ASC the way it does in NT populations. Decades of research into the neurobiological basis of imitation has identified several key networks that help delineate the social bonding function of imitation from its learning function (Hamilton, 2015; Iacoboni, 2005). According to Hamilton (2015), a core visual-motor stream (comprised of the dorsal and ventral premotor cortex: PMd and PMv, anterior intraparietal sulcus: aIPS, and supramarginal gyrus: SMG) enables sensory integration of information from the environment to produce a matching output. Findings from both behavioural and neuroimaging studies have shown visuo-motor integration impairments specifically associated with imitation deficits in ASC (Haswell, Izawa, Dowell, Mostofsky, & Shadmehr, 2009; Nebel et al., 2016). Yet, there is more to imitation than sensorimotor processing; we do not imitate every action we observe, as selection and control mechanisms are in place that take into account the social factors related to the action observed. Hence, alongside the core stream, the involvement of the medial prefrontal cortex (mPFC), the temporoparietal junction (TPJ), and other frontal regions enables top-down control of context-specific, socially relevant information.

Similarly, Iacoboni (2005) posits a core imitation circuitry that largely corresponds to the so-called mirror neuron system (comprised of the superior temporal sulcus: STS, inferior parietal lobe: IPL, PMd, PMv, and inferior frontal gyrus: IFG). This core circuitry, and its interactions with other regions, is differentially implicated in imitation that serves a procedural skill learning function, as compared to imitation occurring as a form of social mirroring and communication (here referred to as the ‘social bonding’ function).

Accordingly, Iacoboni suggests that the procedural skill learning network is comprised of Brodmann area 46 (BA 46), PMd, pre-supplementary motor area (pre-SMA) and the superior parietal lobe (SPL). In contrast, the social bonding (or mirroring) function of imitation is implicated in interactions of the core circuitry with the insula and the limbic system. Drawing from both Hamilton (2015) and Iacoboni (2005), we examined the neural substrates of mimicry behaviour by contrasting how procedural skill learning versus bonding networks might differentially support mimicry in ASC and NT children.

Prior task-based fMRI studies have shown autism-associated differences in neural activation patterns during social interactions (Di Martino et al., 2009; Mundy, 2018). For instance, it has been found that, while NT adults showed selective activation of frontoparietal regions (i.e., the IFG, premotor complex, precentral gyrus and SMG) in response to social cueing with eye gaze, adults with ASC showed increased activity only in the SPL in response to the social cues (Greene et al., 2011). Notably, this finding maps onto the distinction between the bonding (frontoparietal regions) and procedural skill learning (SPL) regions made above. Using a similar paradigm contrasting social eye gaze cueing with arrow cueing, analogous findings with ASC children have also been reported (Vaidya et al., 2011). While NT children selectively recruited the STS, dorsomedial prefrontal cortex, and middle and inferior frontal regions in response to eye gaze, ASC children showed decreased activity in these areas. These findings are consistent with the possibility that the distinction between the bonding and learning components of social cues may be impaired in ASC, as indicated by atypical representation of stimuli in the learning circuitry. Notably, the suggestion here is not that social learning mechanisms are intact in ASC. Rather, it may be that, in ASC, imitation is predominantly related to learning, and hence, social cues such as mimicry of meaningless gestures are represented in the learning circuitry as well.

Structural MRI studies show widely distributed anatomical variations in the grey matter volume of many brain regions (Foster et al., 2015; Rooij, 2018), including those implicated in social processing (Sato & Uono, 2019). However, vast heterogeneity within the ASC population, and a lack of robust links between neuroanatomical phenotypes and behavioural functions, creates serious obstacles for the observed differences to be used in clinical settings (Ecker, 2017). As a step towards bridging this gap in establishing brain-behaviour associations, in this study, we examined how structural substrates (i.e., grey matter volume) of the procedural skill learning versus bonding regions would be differentially linked to mimicry behaviour in ASC and NT children. We predicted that brain regions involved in social bonding would explain mimicry frequency less in the ASC group than in the NT group (Hypothesis 3). Further, we hypothesised that, when the contributions of the learning versus bonding regions are contrasted within each diagnostic group, the learning regions would explain mimicry frequency more than bonding regions in the ASC group, while the opposite pattern would be observed for NT children (Hypothesis 4).

To assess mimicry, we used a semi-structured naturalistic interaction task, in which children were observed to see if they spontaneously copied the actions of an interlocutor, who rubbed her head and scratched her arm while telling a story. In addition to this behavioural task, the children completed a structural MRI scan, which was used to obtain the regions of interest within the learning, bonding, and core clusters.

## Results

### Statistical Analysis

For all analyses, the open-source R statistics software has been used(R Core Team, 2013). Mimicry frequency was assessed by calculating the instances of mimicry based on the criteria detailed in the Methods section. The mimicry outcome variable was highly positively skewed, not meeting the normal distribution assumptions of a general linear model. Hence, we used the square root of the outcome variable in all future analyses. To examine the effect of diagnosis on mimicry, we also used a non-parametric Kruskal-Wallis test on the raw mimicry frequency data and plotted those raw values in Figure 1.

**Figure 1.**
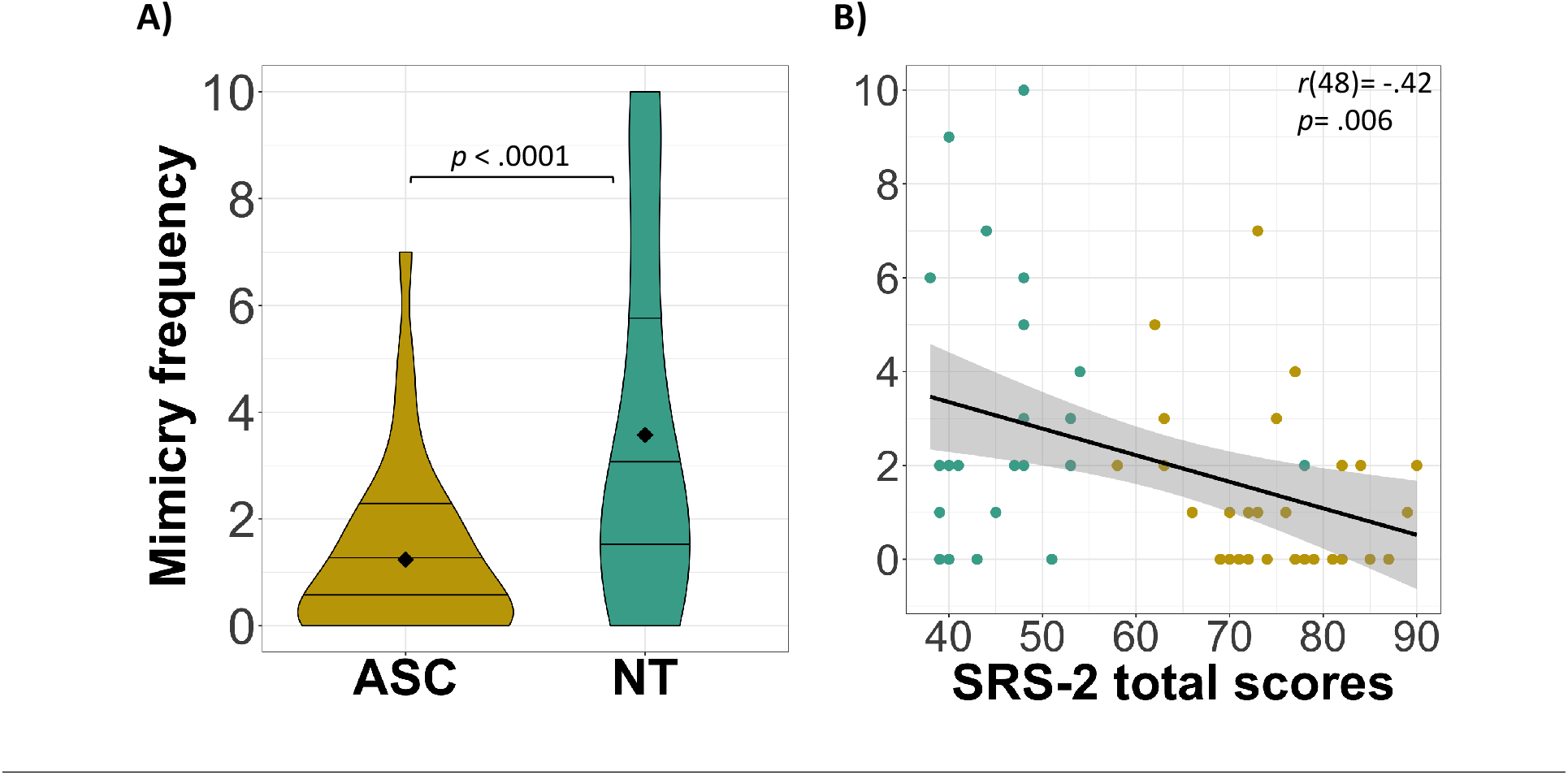
Mimicry and autism diagnosis. **A)** Violin plots showing the difference in the frequency of mimicry between the ASC group and the NT group. Black diamonds indicate the mean and horizontal lines indicate the 25%, 50% and 75% quartiles. **B)** Scatter plot showing the association of mimicry frequency with autism severity as assessed using SRS-2 total T-scores.

To examine how mimicry was associated with brain structures, the grey matter volumes of our regions of interest were calculated (see Methods for details). The regions of interest were selected based on the works of Iacoboni (2005) and Hamilton (2015), as well as the literature reviewed above. The *procedural skill learning regions* were comprised of the dorsolateral prefrontal cortex (DLPFC; corresponding to Iacoboni’s BA46), ventral premotor cortex (PMv; corresponding to Iacoboni’s Pmd), supplementary motor complex (corresponding to Iacoboni’s pre-SMA) and the superior parietal lobule (SPL). The *bonding regions* were comprised of the insula, the limbic system (i.e., amygdala, thalamus, hippocampus) and medial prefrontal cortex (mPFC). A core visuo-motor cluster was also defined as being comprised of the inferior parietal lobe (IPL), superior temporal sulcus (STS) and supramarginal gyrus (SMG). To examine the relative contributions of different brain regions for mimicry within each diagnostic group (ASC vs NT), we conducted robust bootstrapping simulations using the relaimpo package in R (Groemping & Lehrkamp, 2018).

#### 1. Mimicry frequency and autism

First, we examined whether the ASC and NT groups exhibited different amounts of mimicry (Hypothesis 1). A linear regression model with diagnosis (ASC vs NT) as the predictor variable, age and sex as covariates, and the square root of mimicry frequency as the outcome variable, confirmed our hypothesis by showing that children with ASC (*M*= 1.24, SD= 1.63; see Figure 1A) mimicked the story-teller significantly less than did NT children (*M*= 3.57, SD= 3.44; *β*= 0.83, SE= 0.24, *p*= .001). A non-parametric Kruskal-Wallis test examining the effect of diagnosis (ASC vs NT) on the raw mimicry frequency values similarly showed significantly reduced mimicry in ASC as compared to NT (*X^2^*(1)= 9.21, *p*= .001).

To examine Hypothesis 2, we assessed how mimicry frequency (in raw values) was correlated with the severity of autism symptoms as assessed by the SRS-2 (administered to the entire sample) and ADOS-2 (administered to the ASC group only). This analysis revealed that, across diagnostic groups, increased mimicry was strongly associated with decreased autism symptoms as indicated by the SRS-2 total T-scores (*r*(48)= −.42, *p*= .006; see Figure 1B). The association between mimicry and ADOS-2 total scores did not reach significance, presumably due to small sample size (i.e., n= 34 for the ASC group) and restricted variance (*p*> .05; see Table 1). This finding both supports our hypothesis and demonstrates the clinical relevance of spontaneous mimicry as a behavioural indicator of autism severity.

**Table 1.** Bivariate Pearson correlations of mimicry, bonding, learning and core visuo-motor clusters with core autism symptoms as measured by SRS-2 and ADOS-2 scales, whereby higher scores indicate increased autism severity (ASC, NT, **overall** across diagnostic groups)*. P values are corrected for multiple comparisons.* * *p*< .05, ** *p*< .01.

|  | Mimicry | Bonding cluster | Learning cluster | Core v-m cluster |
| --- | --- | --- | --- | --- |
|  | -.42** | .11 | .08 | .20 |
| SRS-2 total | -.24 | -.12 | -.06 | -.34 |
|  | .17 | -.08 | -.19 | .0006 |
| ADOS-2 total | -.28 | -.43* | -.30 | -.50* |

#### 2. Diagnostic differences in the relationship between mimicry and brain regions

To examine Hypothesis 3 – that mimicry is linked to social bonding less in ASC than in NT – we conducted three multiple linear regression tests, one per brain cluster (5000 bootstraps). In these models, diagnosis (ASC vs NT) was the moderator variable, total grey matter volumes of the learning, bonding, and core clusters were predictor variables, and the square root of mimicry frequency values was the outcome variable. All tests were covaried for participant age, gender, and whole-brain grey matter volume. The interaction of diagnosis with the bonding regions was significant (*β*= .0003, *SE*= .0001, *p*= .01, R^2^= .32), such that larger grey matter volume of the bonding regions was associated with mimicry frequency more in the NT group (R^2^= .37) than in the ASC group (R^2^= .12). In contrast, the interaction term was insignificant for the learning (*β*= .00006, *SE*= .00007, *p*= .39, R^2^= .21) and core (*β*= .00005, *SE*= .00008, *p*= .53, R^2^= .16) clusters. This finding shows that, as hypothesised, the bonding regions contributed to mimicry significantly less in ASC than in NT, while the learning and core clusters contributed similarly to mimicry in both ASC and NT groups.

Lastly, we investigated the relative contributions of each brain region (learning cluster: DLPFC, supplementary motor cortex, SPL, PMv; bonding cluster: mPFC, limbic system, insula) towards explaining mimicry within the diagnostic groups (Hypothesis 4). To calculate the R^2^ contributions of each region, 5000 simulations were done on the relaimpo package by setting the type parameter to lmg, which represents the R^2^ value partitioned by averaging over orders. In the multiple regression models, brain regions of the learning and bonding clusters were the predictor variables, and the degree of mimicry was the outcome variable. All models were controlled for total cerebral volume (TCV). The relative contributions of each cluster and region per diagnosis can be seen in Figure 2.

**Figure 2.**
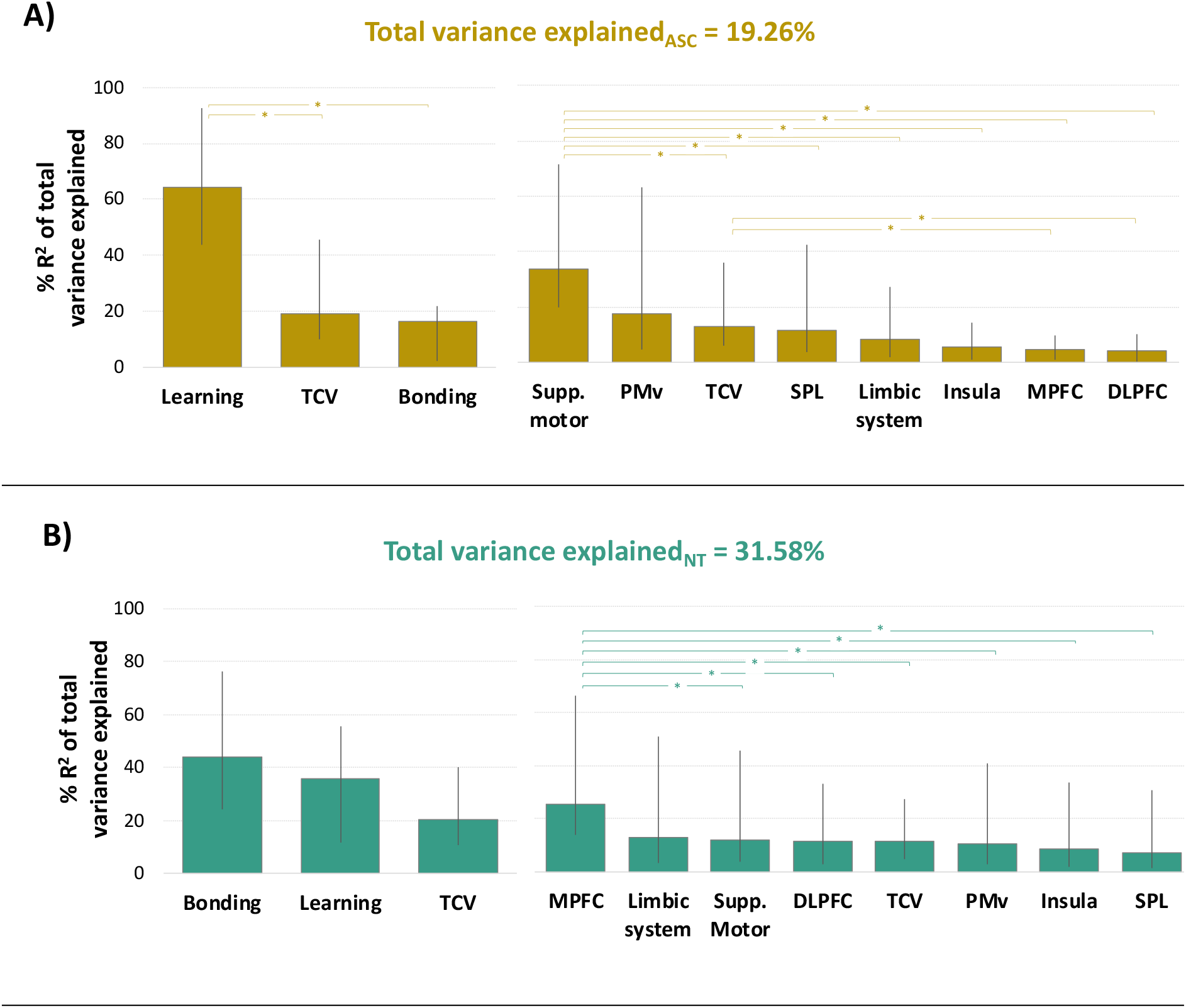
Relative contributions of brain regions for explaining the variance in the amount of mimicry **A)** for the ASC group and **B)** for the NT group. Bars indicate 95% bootstrap confidence intervals and stars indicate significant pairwise differences in contributing regions.

Within the ASC group, the seven brain regions explained 19.26% of the variance in mimicry, with the learning cluster explaining a significantly greater proportion of this variance (64.34%) as compared to both the bonding cluster (16.43%) and the whole brain volume (19.22%), which were not different from each other (CIs for learning vs bonding: [−.919, −.214]; learning vs TCV: [.095, .808]; bonding vs TCV: [−.289, .013]). Examination of the brain regions individually revealed that the largest contributor was the supplementary motor cortex (33.74% of all variance explained), which was significantly different from the contributions of all other regions except for PMv (see Table S2 for test statistics).

Similar bootstrapping simulations in the NT group revealed that the seven brain regions explained a total of 31.58% of the variance in mimicry, with the bonding cluster having the largest contribution (43.80%), which was not significantly different from the contributions of the learning cluster (35.90%) or the whole brain volume (20.30%), with the latter two also not being significantly different from each other (CIs for learning vs bonding: [−.307, .636]; learning vs TCV: [−.192, .366]; bonding vs TCV: [−.006, .621]). When the regions were examined individually, the largest contributor was the mPFC (25.36% of all variance explained), which was significantly different from all other regions except for the limbic system and TCV (see Table S2 for test statistics).

The findings on relative contributions of individual brain regions with each group (ASC and NT) support our hypothesis that, while mimicry is explained less by bonding and more by learning regions within the ASC group, the reverse trend is observed within NT.

As an exploratory analysis, we also examined the bivariate correlations between our brain regions of interest (i.e., the bonding, learning and core visuo-motor clusters) and autism severity. The grey matter volumes of regions comprising the bonding cluster (*r*(22)= −.43, *p*= .04) and the regions comprising the core visuo-motor cluster (*r*(22)= −.50, *p*= .02) were negatively associated with ADOS-2 total scores; no brain region significantly correlated with the SRS-2 scores. In relation to our hypotheses about the procedural learning versus bonding functions of mimicry, this finding implies that differences in the structural properties of the bonding cluster, rather than the learning cluster, are more closely associated with autism symptoms.

## Discussion

This study investigated how spontaneous mimicry of gestures during an interaction relates to learning and social bonding in children with and without ASC. We found that children with ASC mimicked their interaction partners less than did NT children, and that the amount of mimicry was negatively associated with autism symptom severity. The examination of brain-behaviour associations further revealed that structural properties (i.e., grey matter volume) of brain regions involved in *social bonding* explained mimicry less in children with ASC than in NT. Mimicry in ASC was better explained by brain regions involved in *procedural skill learning*.

Do individuals with ASC not copy others’ actions as frequently, and if so, why not? This has been a question of debate for decades (Edwards, 2014; Williams, Whiten, & Singh, 2004). Previous research has established autism-associated deficits in copying others’ actions, especially when the actions seem meaningless or not goal-oriented (Eigsti, 2013). Similarly, brain networks involved in imitation seem to be activated differently between ASC and NT populations (Hamilton, 2015; Iacoboni, 2005). Yet, a crucial distinction, namely the distinction between procedural skill learning and social bonding functions of imitation, has been largely overlooked in this discussion. To understand more about why mimicry occurs less frequently in ASC and what, if any, implications this has for the social-communicative impairments observed in autism, it is essential to determine whether or not copying actions is linked to social bonding in ASC.

Our behavioural data reveal that children with ASC did indeed copy their interaction partner’s gestures less frequently than did their NT peers. This finding adds to a growing body of research, which has showed altered mimicry of facial expressions presented to ASC participants on computer screens (Helt et al., 2020; Oberman et al., 2009; Senju et al., 2007) and of gestures performed during live interactions (Helt et al., 2010). Importantly, our interaction task was designed to be naturalistic, and yet, not biased towards NT interactions, which are characterised by coordinated exchanges such as mutual eye gaze and turn-taking (Akhtar & Jaswal, 2020). A naturalistic environment was attained for both ASC and NT participants by making the video as lifelike as possible and introducing the interaction task as a semi-structured memory game, while any potential experimenter bias and performance demands were eliminated through presenting the partner’s actions via a pre-recorded video. Furthermore, we demonstrate the clinical relevance of mimicry with the finding that mimicry frequency was negatively associated with autism symptom severity across all children (as measured by SRS-2). These findings thus suggest that reduced mimicry may in part explain the social-communicative difficulties commonly observed in ASC.

Beyond demonstrating diagnostic differences in the amount of mimicry, this study shows how mimicry was associated with brain regions involved in procedural learning versus social bonding. Addressing the potential problem of spurious findings in neuroscience research (Vul, Harris, Winkielman, & Pashler, 2009), we used a strictly theory- and hypothesis-driven approach for defining our brain regions of interest a priori. As predicted, the association between mimicry frequency and grey matter volumes of the social bonding regions was significantly weaker in ASC than in NT children. Previous research has shown that deficits in instructed imitation are associated with altered on-line activation of areas comprising our social bonding regions (Di Martino et al., 2009; Mundy, 2018). Our findings extend this prior work by showing that mimicry deficits in ASC can be characterised by their weaker association with the structural properties of the social bonding regions. Linking a behavioural deficit with brain structural properties may indicate a fundamental developmental difference, which calls for examination of this relationship at younger ages.

Finally, we explored how the procedural learning versus social bonding regions contributed to mimicry behaviour within the ASC and NT groups. This analysis showed that, while mimicry was best explained by procedural learning regions in ASC, the largest contributor in NT children was social bonding regions. More specifically, we found that the greatest contributors to mimicry were the supplementary motor cortex and ventral premotor cortex in ASC, and the medial prefrontal cortex (mPFC) and the limbic system in NT children. The mPFC plays an important role in imitation by modulating the sensory input that carries social information, such as eye gaze (Wang, Ramsey, & Antonia, 2011). The limbic system has been implicated in the observation and imitation of emotional facial expressions (Carr, Iacoboni, Dubeau, Mazziotta, & Lenzi, 2003). Therefore, a disassociation of the mPFC and the limbic system with mimicry behaviour may mean that the gaze and facial cues conveyed through mimicry are not robustly perceived in individuals with ASC.

Notably, if mimicry is dissociated from social bonding homogenously within ASC populations, this would have important implications for how ‘social-communicative impairments’ are defined in autism. Previous research has shown that despite altered (and often reduced) imitation by individuals with ASC (Eigsti, 2013), being imitated by others can nevertheless help improve social skills in children with ASC, especially those with low developmental level (Contaldo et al., 2016). This suggests that observed mimicry deficits in ASC are not as innate as the ‘broken mirror neuron’ hypothesis claims them to be (Southgate & Hamilton, 2008). Instead, a view gaining increasing traction is that the seemingly abnormal social interactions observed in ASC may stem from a ‘double-empathy’ problem (Milton, 2012). According to the ‘double-empathy’ problem, individuals with different ways of perceiving and experiencing the world around them would have difficulty communicating with each other. In support of this approach, a growing line of research is showing how ASC-ASC interactions are smoother and more satisfactory for the interacting partners than ASC-NT interactions (Crompton, Hallett, Ropar, Flynn, & Fletcher-Watson, 2020; Sinclair, 2010). These findings put into question whether an observed behavioural difference in ASC as compared to NT should be viewed as an ‘impairment’ and, if so, under which conditions. In the context of mimicry, it is therefore crucial for future research to examine whether reduced mimicry impedes social interactions in ASC-ASC interactions as well, or whether the effect is unique to ASC-NT interactions.

Several limitations of this research should be considered while evaluating the findings. Firstly, our sample included children above a certain IQ threshold. The generalisability of our findings is therefore restricted to children who are not highly intellectually or verbally impaired. Another possible limitation of this study is the correlational nature of the measures taken, as we rely on natural variability in the structural properties of the brain. To provide a ‘baseline’, future research can examine these questions with longitudinal methods. Additionally, it would be insightful to explore these links using behavioural and/or psychophysical measures of bonding, such as markers of the endogenous opioid system and/or the oxytocin/vasopressin system (MacHin & Dunbar, 2011), applied simultaneously with the mimicry task to supplement our neural findings.

Copying others’ actions is a universal feature of social interactions in individuals with neurotypical development, and is known to serve the functions of procedural skill learning and forging social bonds. Our findings shed light on this crucial distinction between the two functions by showing that spontaneous mimicry of actions not only occurs more rarely in ASC, but also that, when it occurs, it is less linked to social bonding and more to procedural skill learning. A dissociation between mimicry and social bonding domains may partly explain the social-communicative impairments that characterise ASC.

## Methods

### Participants

A thorough set of inclusion and exclusion criteria were used to accurately characterise the ASC and NT groups. For all children in the ASC group, autism diagnosis was confirmed by a child neurologist (senior author S.H.M.) according to DSM-5 criteria, using the ADOS-2 module-3 and the Autism Diagnostic Interview-Revised (ADI-R). Children were included in the study only if they had a minimum score of 80 for Full-Scale IQ or one of the indices (Verbal Comprehension, Visual-Spatial, or Fluid Reasoning) on the Wechsler Intelligence Scale for Children, Fifth Edition (WISC-V). For all children, parent-report of Social Responsiveness Scale (SRS-2) was obtained. NT children were eligible if they did not have any diagnosed developmental or psychiatric disorder. Full details of the inclusion/exclusion criteria and characterisation of the diagnostic groups can be found in Table S1 in Supplementary Information (SI).

A total of 74 children (28 NT, 46 ASC), aged 8-12, participated in the mimicry study. Out of this sample, 4 NT and 8 ASC children did not have valid mimicry data due to experimenter error (i.e., misplacement of the camera and data loss: 4 NT, 6 ASC), children’s inattentiveness (3 ASC), and one ASC child not meeting the diagnostic criteria. Thus, the final sample used in the mimicry analyses was comprised of 60 children (**NT group**: n= 28, 3 girls, M_age_= 10.23, M_IQ_= 110.24; **ASC group**: n= 32, 1 girl, M_age_= 10.38; M_IQ_= 103.19). An additional 11 children (4 NT, 7 ASC) were excluded from imaging analyses (n= 5 due to not having been scanned, n= 6 due to having poor data quality). The neuroimaging analyses therefore included 24 NT and 24 ASC children (**NT group**: 2 girls, M_age_= 10.13, M_IQ_= 111.05; **ASC group**: 1 girl, M_age_= 10.21, M_IQ_= 104.57). In both datasets, the ASC and NT groups were balanced on IQ as assessed by the WISC-V (both *p*’s> .05; see Table S1).

This study was approved by the Johns Hopkins School of Medicine’s Institutional Review Board. All participants’ legal guardians provided written consent prior to the study, and child participants provided written assent. The study took place as part of two day-long visits to the Center for Neurodevelopmental & Imaging Research at the Kennedy Krieger Institute; participating families were compensated $100 for their time.

### Procedure

#### Mimicry task

Mimicry was assessed during a story-telling task, in which the children first watched a video of a woman telling them a story, and then were asked to narrate the story back to her. To increase engagement and decrease memory load, the story was split into 5 blocks. Each block lasted for about 2.5 minutes, with unlimited time in-between for children to retell that part of the story. The first 2 blocks were “baseline” blocks, meant to assess the children’s natural tendency to gesture; during baseline, the narrator did not perform any target action. However, in the last 3 blocks, the narrator rubbed her face and scratched her arm (once each per block). The number of times the children rubbed their face or scratched their arm following the narrator’s action within the same block was used as the measure of mimicry frequency. To calculate mimicry scores, the frequency of target actions performed spontaneously at baseline (if any) was deducted from the frequency of target actions performed in the last 3 blocks after the narrator had performed them.

Several procedural measures were taken against potential social interaction pressures and biases. Video recording was used, rather than live interaction, to control for inadvertent social cues from the narrator, such as eye contact or smiles, which could influence participants affiliating with and mimicking the narrator. Still, to attain a naturalistic interaction environment, the interlocutor was never referred to as a “video” and she nodded and seemed to be listening to the child while they retold the story back to her. Moreover, the task was presented to the participants as a ‘memory game’, to prevent the children from focussing on imitation as the purpose of the task. In the video, which was presented on a large TV screen, the narrator was seated in the same room setup as the child. At the end of the session, we asked the children if they noticed “anything unusual” and then asked whether they noticed the narrator’s face rubbing or arm scratching. If children reported noticing either action, then incidents of that action were excluded from the child’s mimicry score. Four ASC children noticed arm scratching, and one NT child noticed face rubbing; out of these, only one ASC child copied an instance of arm scratching, which was removed from further analysis.

#### MRI scan preparation

The children were instructed to relax and watch a movie or take a nap while remaining still for the duration of the scan, which lasted approximately 6 minutes. All children completed a mock scan training session prior to the actual scanning session during which a behavioural protocol designed for children with developmental disabilities was employed. The mock scan aimed to familiarise the children with the scanner environment (e.g., scanner noises, movement, and bore) and to reduce head and body movement during scan acquisition. The behavioral procedures employed have been used and described in detail in previous research (Mahone et al., 2011).

#### Structural MRI acquisition

High resolution T1-weighted Magnetization Prepared Rapid Gradient Recalled Echo (MPRAGE) images were acquired of the entire brain on a Philips 3T MRI scanner (Best, the Netherlands) using a 32 channel head coil (repetition time = 8.0 ms, echo time = 3.7 ms, Flip angle = 8°, voxel size = 1 mm isotropic). A Sensitivity Encoding (SENSE) coil was used to address geometric distortion artifacts due to macroscopic magnetic susceptibility effects that can cause signal dropout at the air-tissue interface.

To ensure good data quality, excessive motion was assessed in 3 steps: First, MPRAGEs were visually inspected at the scanner for gross motion artifacts (e.g., ringing, pixelation, frame shifts, blurred gray-white matter boundaries), and new MPRAGEs were collected if excessive motion was detected and corrective feedback provided to the child). Second, the MPRAGE quality was rated on a 5-point scale (Good, Borderline +, Borderline, Borderline-, Unusable), which reflected the amount of motion artifact detected by a trained research team member blind to the participant’s clinical diagnosis. Third, regional segmentation maps were visually inspected for errors. Scans with a rating of Borderline- or Unusable were excluded from these analyses (n= 6).

### Data processing

#### Mimicry data processing

All children were video-recorded during the entire session for post-hoc coding purposes. Using the open-source E-LAN software version 5.7 (2019), the videos were coded by annotating the start- and end-points of all observed face rubbing and arm scratching actions as performed by the children. Whilst coding for the child’s actions, the raters were naïve to the timing of the narrator’s actions. Once the raters finished coding for the child’s actions, the video was unmuted and viewed again to annotate the timestamps of the blocks and the narrator’s target actions. The E-LAN output with the timestamps was processed using a custom-written script on Matlab 2018b. This script calculated the timestamps of only those target actions of the children that occurred after the narrator had performed the same action within that block. Additional variables, such as the duration of children’s retelling and latency of mimicry, were also extracted, though are not reported here.

#### Structural MRI data processing

The limbic regions, including amygdala, thalamus and hippocampus, were derived from MR images using a validated hierarchical segmentation pipeline (Tang et al., 2015), which is built upon a two-level multi-atlas likelihood-fusion (MALF) algorithm in the framework of the random deformable template model (Tang et al., 2013). The pipeline’s reliability and accuracy in segmenting the subcortical structures from MR images has been established in a previous study (Tang et al., 2015), and all segmentations were visually inspected for accuracy. Cortical grey matter volumes were derived using FreeSurfer (Fischl et al, 2004). Within FreeSurfer, the Ranta atlas (Ranta et al, 2014) was used to delineate the regions: DLPFC, PMv, supplementary motor complex and mPFC, and the Desikan atlas (Desikan et al, 2006) was used to delineate the regions: SPL, insula, IPL, STS and SMG.

We derived the regions of interest based on the core visuo-motor, learning and social bonding networks indicated in the previous literature. The <u>core visuo-motor cluster</u> was comprised of the inferior parietal lobe (IPL), superior temporal sulcus (STS) and supramarginal gyrus (SMG). Following the distinctions made between bonding and learning functions in the Iacoboni (2005) and Hamilton (2015) models, the <u>bonding cluster</u> was comprised of: the insula, limbic system and medial prefrontal cortex (mPFC). The temporoparietal junction suggested by Hamilton was left out of our bonding cluster, because our atlases did not distinguish this specific region. Finally, the <u>learning cluster</u> comprised of all the regions suggested by Iacoboni’s model, namely: the dorsolateral prefrontal cortex (DLPFC; corresponding to Iacoboni’s BA46), ventral premotor cortex (PMv; corresponding to Iacoboni’s Pmd), supplementary motor complex (corresponding to Iacoboni’s pre-SMA) and the superior parietal lobule (SPL).

Given that our hypotheses were not specific to either hemisphere, after confirming that the left and right hemispheric grey matter volumes were highly correlated for each brain region of interest (all *p*s< .0001), we summed them up to create a single grey matter volume value per region.

## Supplementary Information

### Participant inclusion/exclusion criteria

Participants were screened to exclude individuals with co-occurring neurological or medical conditions that might confound the results including (i) known genetic disorder (e.g., NF1, tuberous sclerosis), acquired neurologic disease (e.g., stroke, tumour), cerebral palsy, history of severe head injury, intracranial pathology or significant dysmorphology, (ii) history of seizures or confirmed diagnosis of epilepsy, (iii) any progressive (e.g., neurodegenerative) neurological disorder, (iv) history of head injury resulting in prolonged loss of consciousness, (v) active psychosis, major depression, bipolar disorder, conduct disorder, or adjustment disorder. Presence and history of psychiatric diagnoses were assessed using a comprehensive standardised parent interview, the Kiddie Schedule for Affective Disorders and Schizophrenia for School-Aged Children – Lifetime Version (KSADS). ASC children with co-occurring anxiety and attention deficit hyperactivity disorder (ADHD) were included due to high rates of co-occurrence of these disorders in autism. ASC children on stimulant medication had the medication withheld on the day of testing as well as 24 hours prior.

Children were included in the NT group if they: (1) did not meet clinical criteria for autism spectrum disorders, (2) did not have a history of ADHD, developmental disorder, or other psychiatric disorder based on parent responses from the KSADS, and (3) did not have immediate family members with autism (i.e., a sibling or parent).

## Supplementary results

**Table S1.** Participant characteristics.

|  |  | ASC |  | NT |  | Test statistics<br>(ASC vs NT) |
| --- | --- | --- | --- | --- | --- | --- |
|  |  | Mean (SD) | Range | Mean (SD) | Range |  |
| **Entire**<br>**sample**<br>**(n= 62)** | Chronological age<br>in years | 10.38<br>(1.36) | 8 – 13 | 10.23<br>(1.39) | 8 – 13 | $t(53) = 0.40$<br>$p = .69$ |
| | SRS-2 total score | 74.85<br>(8.08) | 58 – 90 | 46.55<br>(8.68) | 38 – 78 | $t(43) = 12.17$<br>$p < .0001$ |
|  | ADOS-2 total<br>score | 13.94 | 8 – 27 | – | – | – |
| | WISC-V General<br>ability index | 103.19<br>(17.13) | 70 – 148 | 110.24<br>(14.47) | 87 – 141 | $t(55) = -1.79$<br>$p = .08$ |
| | Gender<br>(Boys/Girls) | 33/1 | – | 25/3 | – | $\chi^2(1) = 1.51$<br>$p = .22$ |
| **Sample**<br>**with**<br>**complete**<br>**imaging**<br>**data**<br>**(n= 48)** | Chronological age<br>in years | 10.21<br>(1.35) | 8 – 13 | 10.13<br>(1.33) | 8 – 13 | $t(46) = 0.22$<br>$p = .83$ |
| | SRS-2 total score | 75.65<br>(8.09) | 58 – 89 | 44.63<br>(5.29) | 38 – 54 | $t(38) = 14.93$<br>$p < .0001$ |
|  | ADOS-2 total score | 13.14<br>(3.97) | 8 – 20 | – | – | – |
| | WISC-V General<br>ability index | 104.57<br>(18.06) | 70 – 148 | 111.05<br>(12.36) | 87 – 141 | $t(38) = 14.93$<br>$p = .17$ |
| | Gender<br>(Boys/Girls) | 23/1 | – | 22/2 | – | $\chi^2(1) = 10.35$<br>$p = .56$ |
| | Total cerebral<br>volume (cm<sup>3</sup>) | 1128.28<br>(117.73) | 925.04 –<br>1388.00 | 1079.79<br>(89.76) | 897.47 –<br>1300.24 | $t(43) = 1.60$<br>$p = .12$ |

**Table S2.** Results of the bootstrapping simulations showing pairwise significant differences between regions.

|  | **ASC 95% CI** | **NT 95% CI** |
| --- | --- | --- |
| DLPFC- supp. motor | [-0.652, -0.174] | *n.s.* |
| mPFC – supp. motor | [-0.686, -0.169] | *n.s.* |
| Insula – supp. motor | [-0.687, -0.152] | *n.s.* |
| Limbic – supp. motor | [-0.678, -0.103] | *n.s.* |
| Supp_motor – SPL | [0.056, 0.655] | *n.s.* |
| Supp_motor – whole brain | [0.029, 0.676] | *n.s.* |
| DLPFC – whole brain | [-0.389, -0.017] | *n.s.* |
| mPFC – whole brain | [-0.407, -0.011] | *n.s.* |
| mPFC – SPL | *n.s.* | [0.059, 0.648] |
| mPFC – insula | *n.s.* | [0.039, 0.637] |
| mPFC – PMv | *n.s.* | [0.029, 0.639] |
| mPFC – whole brain | *n.s.* | [0.018, 0.616] |
| mPFC – DLPFC | *n.s.* | [0.023, 0.596] |
| mPFC – supp. motor | *n.s.* | [0.021, 0.618] |

## Notes

### Competing Interest Statement

The authors have declared no competing interest.

